# Positive interactions support complex networks

**DOI:** 10.1101/118166

**Authors:** Gianalberto Losapio, Marcelino de la Cruz, Adrián Escudero, Bernhard Schmid, Christian Schöb

**Affiliations:** Department of Evolutionary Biology and Environmental Studies, University of Zurich, Winterthurerstrasse 190, 8057 Zurich, Switzerland; Departamento de Biología, Geología, Física y Química inorgáanica, Escuela Superior de Ciencias Experimentales y Tecnológicas, Universidad Rey Juan Carlos, C/ Tulipán s/n C.P. 28933 Móstoles (Madrid), Spain

## Abstract

Ecologists have recognised the effects of biotic interactions on the spatial distribution of living organisms. Yet, the spatial structure of plant interaction networks in real-world ecosystems has remained elusive so far. Using spatial pattern and network analyses, we found that alpine plant communities are organised in spatially variable and complex networks. Specifically, the cohesiveness of complex networks is promoted by short-distance positive plant interactions. At fine spatial scale, where positive mutual interactions prevailed, networks were characterised by a large connected component. With increasing scale, when negative interactions took over, network architecture became more hierarchical with many detached components that show a network collapse. This study highlights the crucial role of positive interactions for maintaining species diversity and the resistance of communities in the face of environmental perturbations.

---

The nature of biodiversity continues to intrigue biologists because of the complexity of interactions among species in ecosystems. Standard ecological theory assumes that negative interactions between species such as competition are essential to promote stable species coexistence^1,2,3,4^. However, recent studies emphasised the importance of positive interactions such as mutualism and facilitation for biodiversity maintenance and ecosystem functioning^5,6,7,8^. Particularly, an impressive amount of studies about networks of mutualistic interactions between plants and animals has increased our understanding of ecological and evolutionary processes shaping communities and ecosystems^9,10^. Conversely, networks of interactions among plants have been less explored. Nevertheless, the existence of interaction networks among multiple plant species has been recently revealed using models of intransitive competition in fully-connected graphs^11,3,12, facilitation by keystone species in bipartite networks^13,14^ and fine scale co-occurrence models for unipartite networks^15,16^^.

Biotic interactions can have consequences on the distribution of organisms and shape the spatial structure of populations and communities. Specifically, competitive interactions can promote fine-scale segregation^17,1,18,19^, while facilitative interactions can promote fine-scale aggregation^5,20,21,22^. Consequently, if microhabitat conditions and stochasticity are taken into account it is possible to consider fine-scale spatial aggregation (i.e. significantly positive associations) and spatial segregation (i.e. significantly negative associations) as indicators of facilitation and competition, respectively. Analogously, non significant spatial dependency can indicate neutral net interactions. By considering spatially explicit models, recent studies suggest that the outcome of positive plant interactions may be diffuse, involving many species^22^ and varying with spatial scale^19^. Furthermore, increasing evidence highlights the importance of indirect interactions for structuring plant communities^23,24,25,26,27^. However, little is known about how plant–plant networks are structured across spatial scales and which network-level factors could maintain species diversity. Directly quantifying the spatial dynamics of plant interaction networks is particularly crucial for understanding how ecosystem processes vary across scales.

To overcome these limitations, we fully mapped a community at the individual-plant level and combined spatial point-pattern with network analyses. We first fitted null models of species distribution and spatial structure for each species. The aim of these null models was to control for niche differences, environmental heterogeneity and stochasticity determining the spatial distribution of each species. Then, we assessed the spatial association among all species to infer species interactions. Although observational approaches are only suggestive regarding the effect of species interactions and other processes, mainly habitat sharing, on spatial association^12^, the observed spatial associations was tested against the expectations of null models of species distribution within the study plot. In this way we accounted for habitat preferences of each individual species. Hence, we assessed wether the observed spatial associations are more or less frequent than expected by hypothetical habitat similarities or differences among species. Finally, we analysed how plant–plant networks changed across spatial scales (Fig. 1) and how they were related to plant richness. Because facilitation is known to be a relevant driver in the examined alpine ecosystem^28,20,29^, we tested the hypothesis that facilitation would support the cohesiveness of plant–plant networks at fine spatial scale, while competition would lead to network disintegration at larger spatial scales.

**Figure 1.**
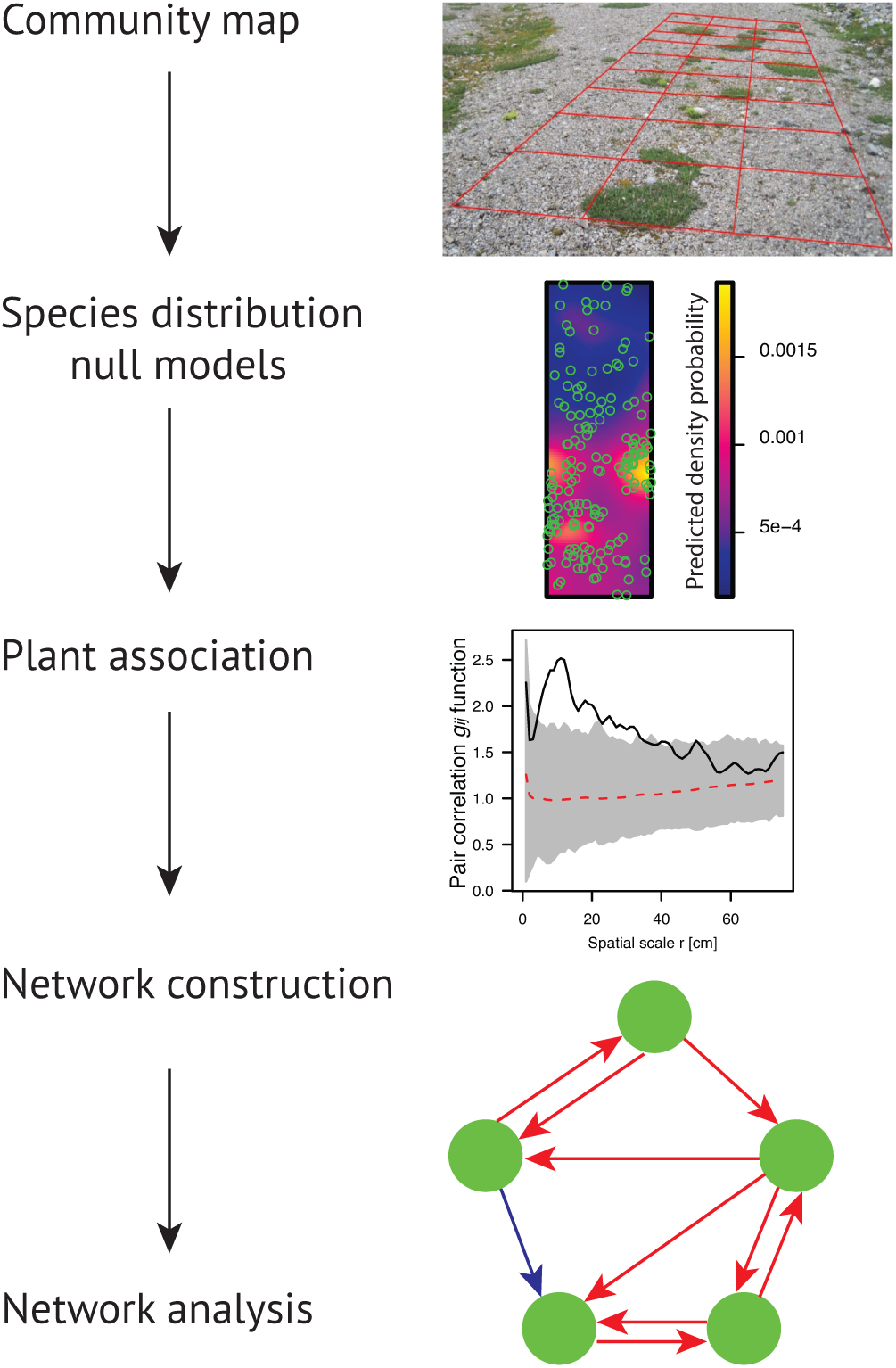
Analytical framework for studying plant interaction networks on the basis of spatial point patterns. A plant community is fully-mapped: for each individual plant, species identity and coordinates are recorded within a spatial grid with a 1 cm accuracy. Spatial point pattern analysis is then employed. First, the distribution of each species is analysed (see Appendix S1 for details). Second, pairwise species associations are estimated after removing the effects of environmental heterogeneity and niche and stochastic processes. Then, species interactions are inferred from spatial association patterns: a positive dependence of species *j* on species *i* is assumed to indicate facilitation of species *i* on species *j*, a negative dependence is assumed to indicate competition, and no association is assumed to indicate neutral interaction. Hence, interaction types are calculated considering the combination between positive, negative and neutral interactions. Finally, network analysis is used to reveal the structural properties, the growth or the collapse of the interaction networks across spatial scales.

## Results

### Shifts of plant–plant interactions across space

A total of 983 interactions were detected across spatial scales among the 19 species. Positive interactions were 592 (60.2%), of which 282 (47.6%) were mutual and 310 (52.4%) were non-mutual. Negative interactions were 391 (39.8%), of which 128 were mutual (32.7%) and 263 were non-mutual (67.3%). No negative–positive interactions were observed. The ratio of positive to negative interactions decreased with increasing spatial scale from 1–75 cm (*β* = −10.294, *β*^2^ = 2.671, *β*^3^ = −2.417, *p* = 0.0001, *R*^2^ = 0.607; Fig. 2a), along with a decrease of the ratio of mutual to non-mutual interactions (*β* = −10.328, *β*^2^ = 6.656, *β*^3^ = 3.606, *p* = 0.0005; *R*^2^ = 0.590; Fig. 2b). This shift from positive to negative interactions went along with a decrease of species richness across spatial scales (Fig. S12). In particular, the richness of interacting plant species increased as the relative amount of positive over negative interactions increased (*β* = 11.798, *β*^2^ = −1.800, *β*^3^ = 4.469, *p* = 0.0019, *R*^2^ = 0.270; Fig. S13).

**Figure 2.**
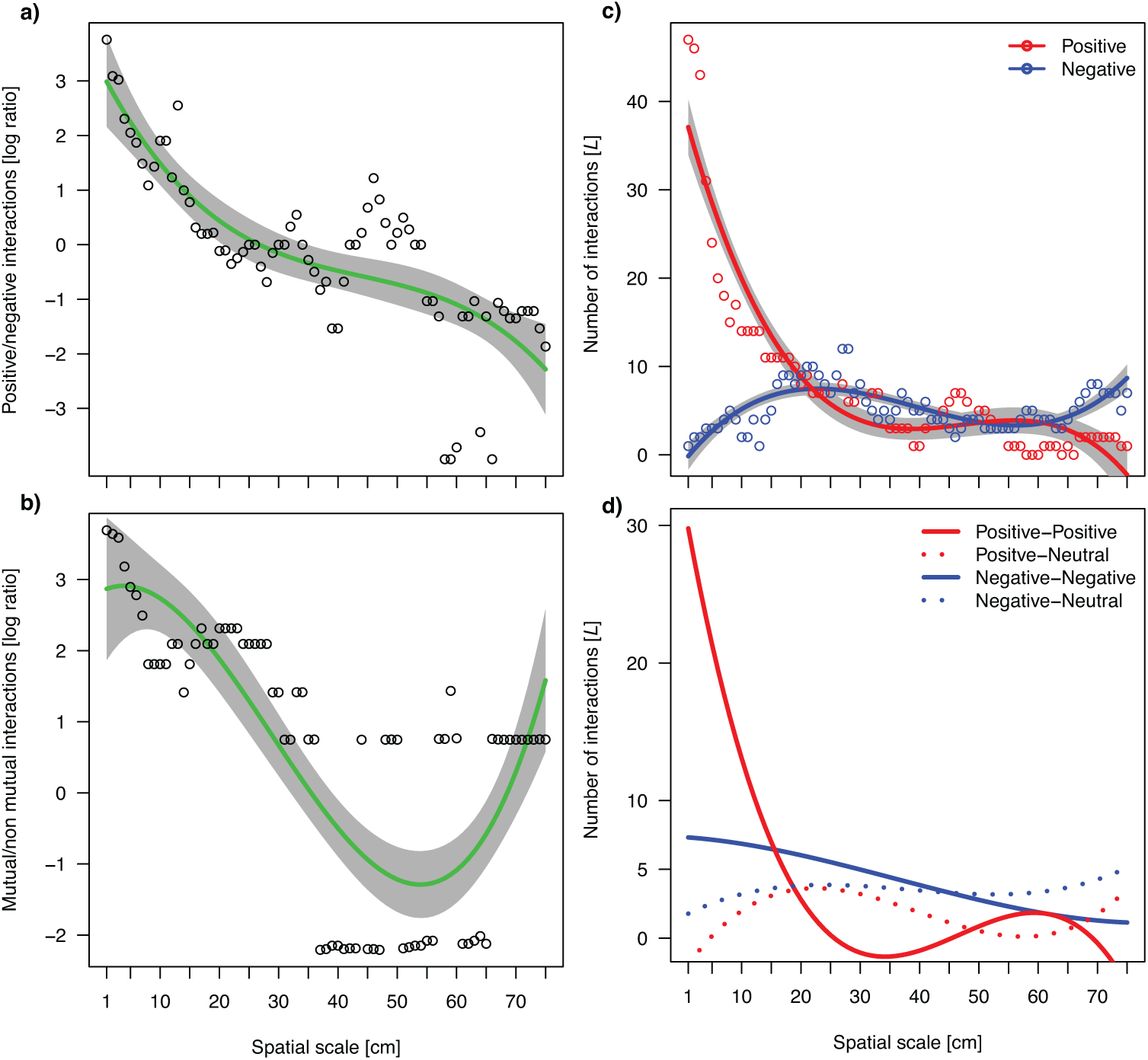
Log ratio between positive and negative interactions **a)**, mutual and non-mutual interactions **b)**, total positive and negative interactions **c)** and total mutual and non-mutual interactions **d)** across spatial scales. Red and blue lines indicate positive and negative interactions, respectively; in **d),** solid and dashed lines indicate mutual and non-mutual interactions, respectively. Predicted lines (i.e. non-linear regression model with the third degree polynomial function of scale as predictor and an autoregressive covariance structure) and 95% CI shown. In **d)** data points and CI omitted for clarity.

### Effects of interaction type

Positive and mutual interactions had a positive effect on the total number of interactions *L* (*p* = 0.0006, *R*^2^ = 0.665; Tab. S3.), while only positive, but not negative, interactions had a positive effect on interacting species richness *S* (*p* = 0.0004, *R*^2^ = 0.630). Thus, there was a decrease in the number of interactions associated with a shift in the predominant interaction type from mutual and positive to non-mutual and negative with increasing spatial scale (*p* = 0.0001, *R*^2^ = 0.607, Fig. 2c-d, Tab. S3).

### Global network architecture

Network clustering gradually decreased within the first 30 cm and then abruptly dropped to 0 with further distance (*β* = −0.970, *β*^2^ = 0.348, *β*^3^ = −0.062, *p* < 0.0001, *R*^2^ = 0.558; Fig. 3a). All interaction-type combinations had significant effects on network clustering (Tab. S4). However, considering their effect size, positive mutual interactions best explained network clustering (*β* = 0.044, *r*^2^ = 0.361, *p* < 0.0001), followed by positive non-mutual interactions (*β* = 0.065, *r*^2^ = 0.225, *p* = 0.0018), whereas negative mutual (*β* = 0.026, *r*^2^ = 0.096, *p* = 0.0247) and non-mutual (*β* = −0.089, *r*^2^ = 0.117, *p* = 0.0139) interactions had weaker effects. This indicates that positive mutual interactions among plants were associated with higher clustering among neighbouring plants.

**Figure 3.**
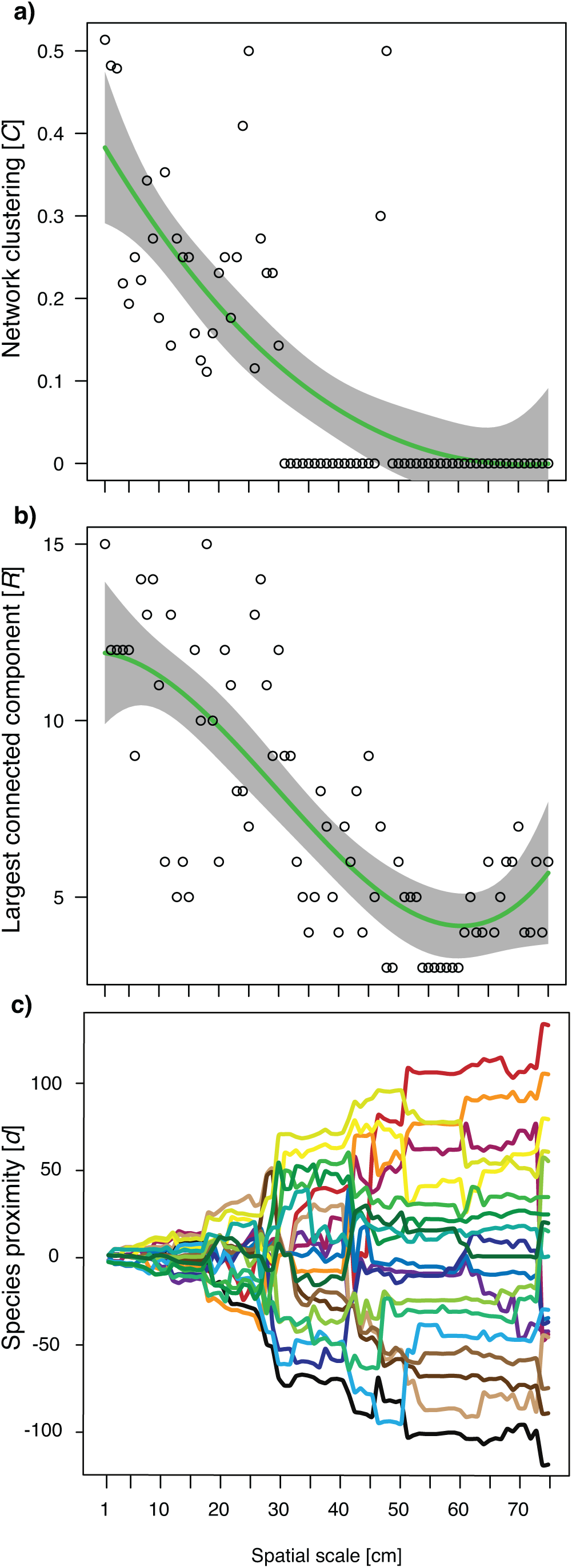
Network transitivity *C* **a)**, size of the largest connected component *R* **b)** and species proximity **c)** across spatial scales. Transitivity, measured by the clustering coefficient *C*37, indicates local cohesiveness of a group of nodes (i.e. species). The size of the largest connected component *R* is the maximum number of interconnected species within a network^44^. A change in the size of the largest connected component provides basic information about the growth of a network. Predicted lines and 95% CI shown. Species proximity calculated on the basis of relative geodesic distance^45^. Each horizontal spline corresponds to a plant species and vertical proximities are proportional to the number of interactions connecting them. The larger the proximity, the higher the fragmentation of the network.

There were connected components across all scales, but their size decreased with increasing scale (*β* = −22.530, *β*^2^ = 6.343, *β*^3^ = 4.270, *p* < 0.0001, *R*^2^ = 0.599) up to about 55 cm (Fig. 3b). Positive mutual and non-mutual interactions and negative non-mutual interactions had significant positive effects on the size of the largest connected component *R* (Tab. S4). Again, positive mutual interactions (*β* = 1.189, *r*^2^ = 0.504, *p* < 0.001) and positive non-mutual interactions (*β* = 2.090, *r*^2^ = 0.383, *p* < 0.0001) best explained variation in *R*, followed by negative non-mutual interactions (*β* = 3.810, *r*^2^ = 0.249, *p* < 0.0001). Species proximity decreased with increasing spatial scale (Fig. 3c). This indicates a network collapse with increasing spatial scale.

## DISCUSSION

Our study highlights the role of positive interactions among plant species for the architecture of complex plant–plant networks. After controlling for niche differences and environmental heterogeneity, we found that facilitation prevailed at spatial scales up to 25 cm, while competition became dominant at spatial scales larger than 50 cm in our alpine ecosystem. This shift from facilitation to competition with increasing distance was coupled with a de-structuring of plant–plant networks, which was ultimately associated with less interacting species. These results support our hypothesis that plant–plant networks change across spatial scales (Fig. 4). Furthermore, they suggest that positive plant interactions could be pivotal in the network organisation of species-rich patches in this stressful, fragmented ecosytem. In summary, at fine spatial scales, positive interactions promoted the cohesiveness of plant–plant networks with high clustering and large connected components. Conversely, at larger spatial scales, networks became more hierarchical and less cohesive in parallel with a relative increase in competitive interactions. Because network complexity may increase ecosystem stability^30^, positive plant interactions may promote plant species richness and ecosystem stability, similarly to obligate plant–animal mutualistic interactions 6.

**Figure 4.**
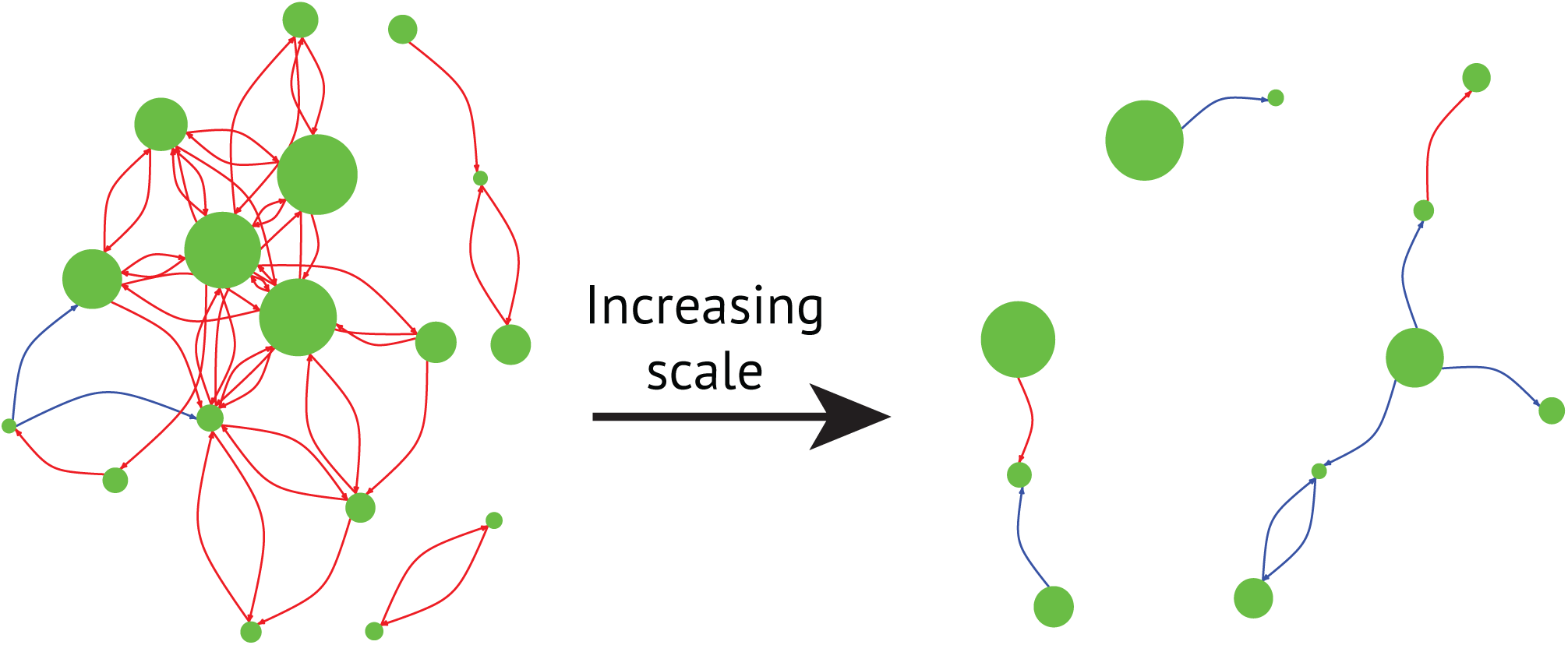
At fine spatial scale (left, 2 cm) positive facilitative interactions (red arrows) build up a network with high transitivity, i.e. high cohesiveness. With increasing scale (right, 50 cm), negative competitive interactions (blue arrows) predominate and the network becomes more disconnected. The size of the nodes (green dots) is proportional to relative species abundance (See Fig. S10 and the online video for the network at every centimetre).

### The spatial scale of plant interactions

Theoretical and empirical studies indicate that the emergence of spatial patterns is due to two main classes of mechanisms of ecological self-organisation^31,30,32,21,33^. The first process considers the role of positive scale-dependent feedbacks between biomass and resources. The second process recognises the role of species as ecosystem engineers and their intraspecific competition. At short distance, plants may increase resource availability, hence ameliorating growth conditions in environments with high abiotic stress as our alpine ecosystem^29,34^. This means that the more plants the stronger the stress amelioration by facilitation can be^21^. Such positive feedback mechanism may explain why facilitation prevailed at the very close proximity to plants, i.e. within vegetation patches. Furthermore, water transport within a patch increases its growth while it inhibits the growth of neighbouring patches. Hence within-patch facilitation may depend on the possibility to exploit resources within and around the patch, thereby leading to between-patch competition^21^. In our case, the importance of competition varied relatively less across scales. Therefore, we suggest that the prevalence of competitive interspecific interactions at larger distances may be associated to resource dynamics between local patches compared to within local patches^1,31,21^. In summary, facilitation may be scale-dependent, whereas competition may be rather constant across space in our fragmented alpine ecosystem.

In addition to these two processes, we postulate here that positive interspecific interactions may be associated with cohesive networks and with the richness of species participating in these networks (Fig. S15). This means that positive interspecific interactions may promote the establishment of more links among different neighbour species. Such effect may result in a facilitation cascade^35^ according to an autocatalytic process^31,30,21^ and similarly to the emergence of cooperation in public goods games^36^. In other words, the presence of positive interactions among neighbouring, diverse plants could be associated with the prevalence of the same positive interactions in the network in plants vicinity. Conversely, at larger distance, the prevalence of negative interactions may reduce the likelihood of species occurring in the network. Ultimately, this may potentially lead to local patches of unexpectedly high species richness characterised by diffuse facilitation^22^.

### The spatial dynamics of plant–plant networks

Networks show a high clustering when the number of interactions among neighbours is large relative to the number of species^37^. The decreasing clustering with increasing scale implies that a transition from a cohesive to a hierarchical organisation of networks occurred in our alpine ecosystem. This shift was nonlinear, but gradual until reaching a threshold at 30 cm, beyond which a sudden, critical transition occurred and clustering rapidly approached zero. This pattern concurs with expectations of the behaviour of an (eco)system approaching a tipping point^30^, highlighting the probable presence of a collapse of the architecture of plant– plant networks. The network collapse could be coupled with the facilitation–competition shift observed across spatial scale in this fragmented system. Potential mechanisms leading to such a shift can be related to previously described positive scale-dependent feedbacks, where positive interactions prevail within patches and negative interactions at larger scale^21^. Coupled to this process there are the positive effects that ecosystem engineers, like *Dryas octopetala* in our system, have on other species^33^, mainly through the decrease of stress and the amelioration of growth conditions 38.

The size of the largest connected components in our networks decreased with increasing spatial scale to half the size at 30 cm and to one-fifth at 55 cm. Again, this reduction in component size was associated with a reduction in positive interactions. In line with this result, we also found a higher species proximity in the network at fine spatial scale where positive interactions were predominant. This indicates that species closely interact at fine scale. On the other hand, species were less closed within the network with increasing scale and negative interactions. Accordingly, the number of cliques (Fig. S14) decreased with increasing spatial scales, indicating network breakdown at its sub-structure level. Taken together, these results suggest a breakdown of the largest connected components with increasing spatial scale, as species tend to segregate into many detached components once positive interactions wane.

Our study is one of the first attempts to analyse the spatial structure of plant–plant networks across scales. We are aware that new questions are now arising. Observational studies such as the present one may suggest potential mechanisms underpinning spatial patterns of species interactions. Nevertheless, with our approach we first controlled for variation in niche differences and environmental heterogeneity before calculating spatial association and then inferring plant–plant interactions^39,19,22^. Moreover, it should be noted that what we observed as facilitation between two species might also be apparent facilitation, in which the two species are both facilitated by a third one. Future experimental studies controlling for differences in demographic stochasticity (e.g. dispersal limitation) and niche processes (e.g. species-specific resource limitation) would be necessary to test the causality of the observed correlations between positive and negative plant–plant interactions with network architecture. At the same time, further theoretical research should accompany such experimental work to better predict network stability under different environmental conditions. We conclude that positive interactions exceed negative ones at fine spatial scales. The resulting increase in network cohesiveness is best supported by the spread of positive interactions among neighbouring plants within the local network in a way that facilitation begets facilitation.

## Methods

### Study area and sampling design

An observational study was performed in a sparsely-vegetated alpine ecosystem (Swiss Alps, 2300 m a.s.l., Lat 46.39995°N, Long 7.58224°E, Fig. S1) characterised by patches of the prostate dwarf-shrub *Dryas octopetala* L. (Rosaceae). The plant community was fully mapped with a 1 cm accuracy during August 2015 within a 9 × 3 m rectangular grid (Fig. S2). For each individual plant (i.e. ramet) we recorded: species identity, coordinates of rooting point (x and y) and a set of functional traits (width, height, number of leaves, leaf dry mass) relevant for resource use and competitive ability^40^. In total, 2154 individuals belonging to 29 species were recorded (Tab. S1). Species richness reached an asymptote in the accumulation curve (Fig. S3), suggesting that a representative area with the entire species pool of this plant community type was sampled. We focused on the 19 species that had more than 10 individuals in order to minimise analytical bias. Fine-scale spatial heterogeneity of soil properties was quantified by determining soil gravel content, soil water content and soil C/N ratio with one composite sample in each 1 m^2^ and beneath each *Dryas* patch (see Appendix S1 for details).

### Spatial pattern analysis and plant interactions

To detect the statistical association between species and infer plant interactions we employed spatial point pattern analysis based on second-order statistics^41,42,39,43^ assuming that spatial patterns could inform about interactions^31,32,30,13,20,21,15^ after accounting for other processes^42,39,43^. The scale of analysis was varied from 1 cm to 75 cm.

First, we describe the spatial distribution of each species. To identify the effects of environmental heterogeneity, niche differences and stochasticity on the species occurrence probability, we fitted different models of spatial distribution within the plot based on species traits, soil properties and stochastic processes for each species. The model with the best goodness of fit was selected as the null model to later test spatial association between species (see Appendix S1 for details).

Second, we determined interspecific spatial associations. We carried out bivariate point pattern analyses for all species pairs to assess the existence of spatial associations between species after accounting for their niche differences and the microenvironmental conditions. We assume that fine-scale spatial segregation and fine-scale spatial aggregation are indicators of competition^17,1,18,19^ and facilitation^5,20,21,22^, respectively. Species association was calculated using the inhomogeneous cross-type pair correlation function *g_ij_*(*r*)39. Given the expected number of points (i.e. individual plants) of species *j* at a distance *r* from an arbitrary point of species *i* (Fig. S4), the probability *p*(*r*) of finding two points *i* and *j* separated by a distance *r* is equal to *p*(*r*) = *λ_i_*(*x*) *λ_j_*(*j*) *g_ij_*(*r*) *dx dy*, where *λ_i_*(*x*) and *λ_j_*(*j*) are the estimated intensity functions of each species (see Tab. S2). Values of *g_ij_*(*r*) > 1 indicate that there are, on average, more individuals of species *j* at a distance *r* from species *i* than expected by chance. Conversely, values of *g_ij_*(*r*) < 1 indicate that species *j* is more segregated from species *i* than expected by chance. When *g_ij_* ≈ 1 the spatial dependency of species *j* on species *i* cannot explain more than what we would expect by chance, i.e. given each species’ distribution.

In order to statistically determine whether an observed pattern was significantly different from what could be expected by chance, Monte Carlo simulation of a realisation of the *g_ij_*(*r*) function at each scale (for *r* from 1−75 cm with 1 cm steps) was used to generate simulated distributions from the null hypothesis of independence of species *j* with respect to species *i*. A total of 199 MC simulations were performed at each scale. The fifth-lowest and the fifth-highest simulated values at each *r* were used to build 95% confidence envelopes around the mean predictions^42,43^. Thus, at a given scale *r*, an empirical 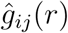 function higher than the confidence envelope indicates significant positive dependence of species *j* on species *i*, while the converse indicates significant negative dependence (Fig. S8, Fig. S9). When 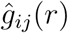 lies within the MC confidence envelope, neutral association cannot be rejected. Because first order constraints on the distributions of each species are controlled (i.e. microsite heterogeneity, niche and stochastic determinants, see Appendix S1), the obtained positive and negative dependences might result from non-random plant–plant interactions^1,31,32,39^. Finally, with this approach we could detect the spatial scales at which such interactions are operating according to the corresponding spatial signals.

### Network analysis

Network analysis was employed to identify the web of plant–plant interactions and to assess how network architecture may promote species coexistence and maintain species richness. At each scale we built a unipartite directed network *G* = (*V, E*) composed of *V* = 19 plant species and *E* ⊆ *V_i_* × *V_j_* significant directional interactions (i.e. distinguishably *E_ij_* and *E_ji_*), for a total of 75 networks and 983 species interactions (Fig. S10 and online video). Each network *G* was represented by an adjacency matrix *M* composed of 19 rows and 19 columns describing interactions among plant species.

Species interactions *E_ij_*(*r*) are described by directed ternary links such that

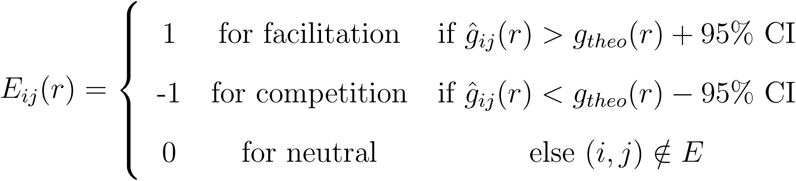

To reveal changes in local plant–plant interactions across scales, for each network we calculated the total number of interactions *E*, the number of species *S* with at least one interaction (*S* < *V*), and the number of pairwise interactions for each bidirectional interaction type, i.e. positive mutual (facilitation–facilitation), positive non-mutual (facilitation–neutral), negative mutual (competition–competition), negative non-mutual (competition–neutral) and negative– positive (facilitation–competition) (Fig. S11).

Network architecture was analysed using the clustering coefficient *C*^37^. *C* tests if two or more species linked to another species are also interacting with each other, measures the local cohesiveness of a group of species and indicates the neighbourhood interaction density as well as the hierarchy and interconnection of a community (Fig. S11). *C* is defined as the probability that neighbouring nodes (i.e. all plant species connected to a plant species *i*) of a plant species *i* are linked to each other. In other words, *C* for any node *i* is the fraction of linked neighbours of *i*, such that 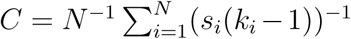, where *s_i_* is the sum of links present among neighbouring nodes for each node *i*, and *k_i_* is the degree (i.e. the number of neighbours) of node *i*. Thus, the higher the clustering, the more the neighbours are connected to each other and the higher the cohesiveness.

To reveal network growth and collapse across spatial scales, we calculated the size of the largest connected component *R.* A connected component of a network is a subset of nodes reachable from every node within it^44^. In other words, the size of *R* is equal to the maximum number of species consecutively linked within a network (Fig. S11). The change in the size of *R* provides basic information about network development and collapse. Hence, the presence of connected components and the change in their size *R* can be used to characterise the robustness of ecological communities.

To reveal network collapse, we calculated species proximity on the basis of relative geodesic distance, i.e. considering nodes positioned on a plane alike^45^. The larger the proximity, the larger the network-based distance among species, the higher the fragmentation of the network.

### Statistical analyses

We first analysed the changes in plant–plant interactions across spatial scales and then we tested the relationships between such changes and network architecture.

We used regression models to relate the response of i) the total number of interactions *E* and ii) the interacting species richness *S* to the ratio between positive and negative interactions, the ratio between mutual and non-mutual interactions and their interactions (fixed effects with third degree polynomials for each ratio, i.e. r + r^2^ + r^3^). Besides, we previously tested with the same approach if the ratio between positive and negative interactions and the ratio between mutual and non-mutual interactions changed across scale (i.e. s + s^2^ + s^3^).

Then, to determine bottom-up effects of local plant–plant interactions on network architecture, we used regression models to test the effects of pairwise interaction combinations (i.e. number of positive–positive, positive–neutral, negative–negative, negative–neutral, negative– positive interactions as fixed effects) on i) the network transitivity *C* and on ii) the size of the largest connected component *R*. By using the absolute number of each interaction-type combination as independent variable we accounted for changes in the total number of interactions across scales. To quantify the importance (i.e. effect size) of the different interaction types and spatial scale, we used the partial r^2^, i.e. the proportion of variation that can be explained by each explanatory variable, calculated as 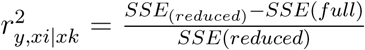, where the error sum of squares SSE (i.e. residuals) were compared between reduced models excluding only one interaction type *x_i_* and the full model containing all interaction types *x_k_*.

We accounted for spatial autocorrelation across scales by including an autoregressive covariance structure (*AR*_(1)_*σ_ij_* = *σ*^2^ *ρ*^|*i*-*j*|^) in all models^46^.

All analyses were done in R 3.3.0^47^, using *spatstat*^43^ and *ecespa*^48^ for spatial pattern analysis, *igraph*^49^ for network analysis and *nmle*^46^ for statistical analysis.

## Data availability

The data that support the findings of this study will be deposited in Dryad repository.

## Acknowledgments

This study was financially supported by the Swiss National Science Foundation (PZ00P3 148261) to CS and partially by the Spanish Ministry of Economy and Competitiveness under the project ROOTS (CGL2015.66809-P) to AE. We thank L. Dutoit and D. Trujillo for their help with data collection in the field. We thank J. Bascompte and M. Fortuna for their fruitful discussions and commenting on an early version of this manuscript.

## Author contributions

GL and CS designed the study, GL collected data and analysed them, MC provided new analytical methods, all authors discussed data analysis, commented the results and edited the manuscript. All authors are included in the author list and agree with its order and they are aware the manuscript has been submitted.

## Materials & Correspondence

Correspondence and requests for materials should be addressed to GL.

## Supplementary Information

Supplementary methods, figures, tables, video, data and R scripts accompany this paper.

